# Testing the accuracy of 3D automatic landmarking via genome-wide association studies

**DOI:** 10.1101/2021.07.22.453315

**Authors:** Yoland Savriama, Diethard Tautz

**Affiliations:** Max-Planck Institute for Evolutionary Biology, Plön, Germany

**Keywords:** Automatic phenotyping, atlas-based segmentation, 3D landmarking, geometric morphometrics, skull, lower jaws, QTL mapping, GWAS, *Mus musculus domesticus*

## Abstract

**Background:** Various advances in 3D automatic phenotyping and particularly in landmark-based geometric morphometric methods have been made, but only a few studies have tested the reliability of such automatic procedures in morphometric analyses. It is generally accepted that automatic landmarking compromises the capture of the actual biological variation, and this not only affects its performance to effectively detect differences among sample means but also the structure of covariance matrices. However, no studies have directly tested the actual impact of such landmarking approaches in analyses requiring a large number of specimens and for which the precision of phenotyping is crucial to capture an actual biological signal adequately.

**Results:** Here, we use a recently developed 3D atlas-based automatic landmarking method to test its accuracy in detecting QTLs associated with craniofacial development of the house mouse skull and lower jaws for a large number of specimens (circa 700) that were previously phenotyped via a semiautomatic landmarking method complemented with manual adjustment. We compare both landmarking methods with univariate and multivariate mapping of the skull and the lower jaws. In the univariate mapping, the automatic approach failed to recover the same SNPs and found only 1 out of 17 previously identified QTLs for the skull, but found one new QTL. Similarly, for the lower jaws, the automatic approach failed to recover the same SNPs but found 2 neighbouring SNPs for 1 out of 8 previously identified QTLs. For centroid size, the same general results were recovered by the automatic method for both the skull and lower jaws, with the same peak SNP being found for the lower jaws. In the multivariate mapping, the automatic approach did not detect the same markers nor QTLs having their regions overlapping with the ones identified with the semi-automatic procedure for the skull, while the same marker, which is also the peak SNP and sole QTL, was recovered by the automatic pipeline for lower jaws.

**Conclusion:** Our results confirm the notion that information is lost in the automated landmarking procedure but somewhat dependent on the analyzed structure. The automatic method seems to capture certain types of structures slightly better, such as lower jaws whose shape is almost entirely summarized by its outline and could be assimilated as a 2D flat object. By contrast, the more apparent 3D features exhibited by a structure such as the skull are not adequately captured by the automatic method. We conclude that using 3D atlas-based automatic landmarking methods requires careful consideration of the experimental question and the cautious interpretation of their results.

## Introduction

Organismal biology is now entering the so-called ‘Era of Big Data’ (e.g., Muñoz and Price 2019) and ‘Phenomics’ (Houle et al. 2010). From the morphological standpoint, this trend reflects the unprecedented progress with analytical devices for the 3D high-definition imaging of morphological phenotypes alongside technological and software development to collect, manage, store, and process the large amount of data produced. One key aspect that must remain central to this revolution in digital morphology is our ability to develop conceptual and methodological frameworks to extract biologically relevant information from these data and address important questions in evolutionary, functional, and developmental morphology. This is where morphometrics, the statistical analysis of shape, plays a major role.

In particular, landmark-based geometric morphometric methods have grown into a mature and powerful branch of biometrics, allowing the capture of morphologically meaningful signals from the diversity and richness of modern imaging data types. These frameworks use a mathematical description of biological structures according to geometric definitions of their size and shape extracted from Cartesian coordinates of points traditionally placed at recognizable structures across specimens (e.g., the intersection of cranial sutures or maximum of bone curvature). Such tasks are particularly highly time-consuming, tedious, and can cause error, especially when substantial sample sizes are involved as in systematics (e.g., Frost et al. 2003) and craniofacial mapping (e.g., Maga et al. 2017; Pallares et al. 2015; Pavlicev et al. 2017; Weiss et al. 2015).

Recently, several atlas-based methods for both automatic segmentation and landmarking have been developed and hold promise for a streamlined treatment of extensive datasets (Devine et al. 2020; Maga et al. 2017; Percival et al. 2019; Porto et al. 2020). These methods use an average volume computed from all available specimens or a subset of them (or a representative specimen), henceforth called an atlas, and apply a suite of image registrations followed by deformations to segment all specimens subsequently. These transformations are re-used to propagate the atlas’ landmark configuration onto all segmented structures (Maga et al. 2017; Percival et al. 2019; Porto et al. 2020) with further refinement that has been suggested to reduce the known systematic difference in means and variance-covariance structure between manual and such automatic procedures (Percival et al. 2019; Porto et al. 2020).

Here, we compare the two approaches by running separate full genome-wide association analyses, which typically require a very high number of specimens and precise phenotyping to identify genes associated with craniofacial development (e.g., Burgio et al. 2009; Katz et al. 2020; Maga et al. 2017; Navarro and Maga 2016; Pallares et al. 2015). To this aim, we re-analyzed a previously published dataset of nearly 700 skulls and lower jaws’ shape and size in an outbred population of male mice acquired via the TINA semi-automatic landmarking tool completed with further manual adjustment (Pallares et al. 2015). The same specimens were processed and automatically landmarked via the Advanced Normalization Tools (ANTs) software (Tustison et al. 2021) and ANTsR (Avants et al. 2015; Tustison et al. 2015) pipelines using a single atlas computed from all of them (Maga et al. 2017; Percival et al. 2019).

In our study, the precision of automatic landmarking is assessed by whether or not (1) the same QTLs identified in the original study are also recovered, (2) the automatic pipeline detects at least the major QTLs, (3) the QTLs discovered by the automatic method have their regions overlapping with the ones that have been initially detected in the original study.

## Material and Methods

### Specimens and data

The specimens analyzed in the present study are the same (circa 700) Carworth Farms White (CFW) outbred mice (Parker et al. 2014) previously analyzed in Pallares et al. (2015), in which these authors used semi-automatic 3D landmark-based geometric morphometrics with additional manual adjustment and high-density genotyping to map genes involved in the craniofacial shape of skulls and lower jaws. All specimens were male and almost the same age (within a two weeks window). Pallares et al. (2015) did not show any significant correlation of age with shape variation, so age was not used as a covariate in the original study.

The genotype data include a filtered set of 80,142 SNPs once genomic markers with a maximum probability genotype above 0.5 and minor allele frequencies below 2% have been excluded prior to the analyses. Morphometric and genomic data were the ones deposited by Pallares et al. (2015) on Dryad (http://dx.doi.org/10.5061/dryad.k543p).

### CT-scanning

Heads in the original study were scanned at a cubic voxel resolution of 21 *μ*m using a *μ*CT Scanco vivaCT 40 (Bruettisellen, Switzerland).

### Landmarks

A configuration of 44 3D landmarks was used for the skull, with 34 placed as mirror images with respect to the midsagittal plane (paired landmarks) and 10 landmarks on top of it (unpaired landmarks). A set of 26 landmarks was used for the lower jaws (13 for each side). See Figure 1 and Table S1 for their definition (same as in Pallares et al. 2015).

**Figure 1.**
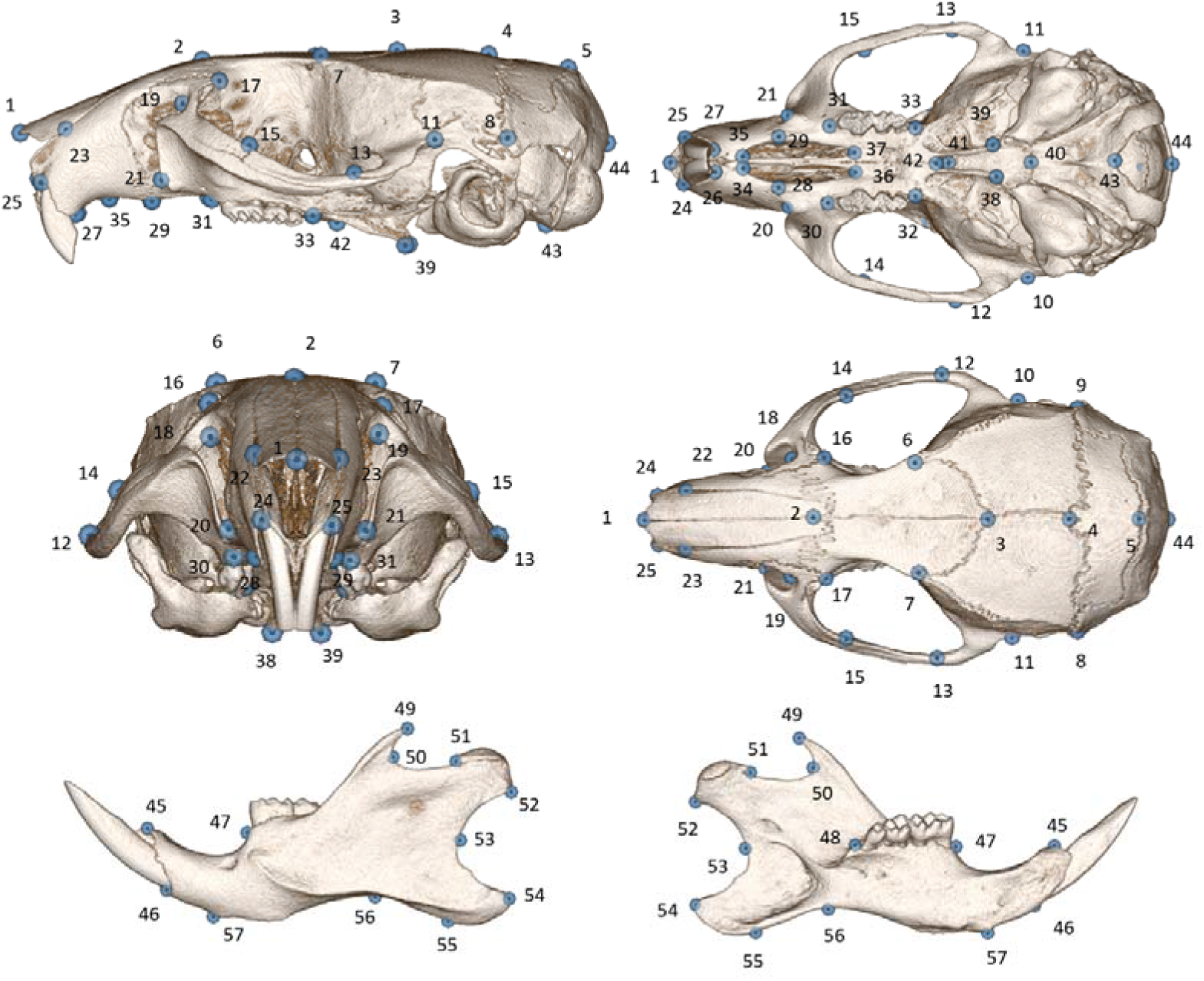
Landmarks used for 3D phenotyping. For the detailed description of the landmarks, see Table S1.

### Semi-automatic landmarking procedure

3D landmark data collection in Pallares et al. (2015) was achieved via a semi-automatic method implemented in the freely available TINA landmarking tool (Bromiley et al. 2014). First, the user computes an average landmark configuration from ten manually landmarked mice. Second, four of these landmarks are manually placed onto each specimen to be used for global registration. Third, a multi-stage registration process using data generated in the previous step is used to propagate the appropriately transformed average reference configuration onto each specimen. The software defines a confidence threshold for each landmark by highlighting them, allowing the user for further manual adjustment whenever needed (Pallares et al. 2015; Pallares et al. 2016).

### Automatic landmarking procedure (ANTs software and ANTsR R package)

Here, we use and extend a recently published full 3D volumetric atlas-based image registration and deformation procedure to automatically segment and generate landmarks on subsequently isolated skulls and lower jaws based on a single manually landmarked average atlas computed from all mice CT-scans (Avants et al. 2009; Avants et al. 2011; Maga et al. 2017; Tustison et al. 2021) https://github.com/muratmaga/mouse_CT_atlas

This procedure uses state-of-the-art medical image registration and a segmentation toolkit implemented in the ANTs software (Avants et al. 2009) https://github.com/ANTsX/ANTs and in its R counterpart ANTsR (Avants et al. 2015; Tustison et al. 2021; Tustison et al. 2015) https://github.com/ANTsX/ANTsR. First, since this pipeline is computationally demanding and typically generates a large amount of data (here 826 GBs), due to the numerous processing steps which output new files each time and especially with such a large sample size (circa 700 CT scans), we reduced the resolution of all scans prior to the analysis. For this purpose, DICOM files of each CT scan were imported in FIJI (Schindelin et al. 2012), converted to 8bits data, downsampled to an isotropic 0.14 mm resolution (same as in Maga et al. 2017) using the Scale option (scaling factor 0.15 with the option ‘Preserve physical image dimensions’ activated and with linear interpolation) in the TransformJ plugin (Meijering et al. 2001), exported as single .nii files using the FIJI NIFTI plugin (Williams 2005), and subsequently compressed using gunzip compression as .nii.gz files. These downsampled files accounted for about 1Mb of file size each so that they could be processed faster in the ANTsR pipeline, while still allowing for the extraction of the main patterns of morphological variation (Maga et al. 2017; Porto et al. 2020).

Second, ANTs was used to compute a single average atlas from all downsampled files generated in the previous step using the script ‘antsMultivariateTemplateConstruction2.sh’ (Avants et al. 2011) https://github.com/ntustison/TemplateBuildingExample. Since specimens were scanned in a standard orientation and consequently were already spatially close to each other, no preliminary registration prior to the template computation was needed to correct for differences in pitch, roll, and yaw, and the default registration steps already included in the ANTs template building script were sufficient. Eight rounds of iteration were satisfactory to generate a template detailed enough to segment the structures of interest for this study, particularly since it was generated from nearly 700 specimens. Here, we chose to generate a single average template since the biological variation was already known to be small; otherwise, specific templates would have been used to accommodate for larger differences. For example, such a registration-based method shows limitations to properly align specimens with a visible size difference compared to the average template. Specimens with observable differences in size should be categorized together, and specific templates should be built for each size category (e.g., Wong et al. 2015; Zamyadi et al. 2010). However, this was not necessary for the samples studied here.

Third, the resulting average atlas was manually segmented using 3DSlicer (Fedorov et al. 2012) http://www.slicer.org, and a labelmap segmenting the skull volume separately from both lower jaws was produced. Since the ANTs and ANTsR image processing both rely on symmetric image deformation and normalization, it was deemed suitable to treat both lower jaws as a single unit that has bilateral symmetry, similarly as for the skull. Tests considering one lower jaw at the time produced unreliable automatic landmarking on such half structure (not shown here). The average template was landmarked using the same 70 landmarks as Pallares et al. (2015) (Figure 1, Table S1) and twice to further test for measurement error.

Fourth, the original approach using ANTsR from Maga et al. (2017) was adapted to extend automatic landmarking to the lower jaws as well. However, the specimens were not registered onto the average template via the function inVariant(), since this function includes an additional rotation step designed for data with highly misaligned specimens and using this function caused misregistration with our data known to contain specimens already close to each other. Instead, the simpler default function antsRegistration() was used here. Prior to segmentation, quick checks were made from 2D slices using function plot.antsImage() to assess each specimen’s overall degree of registration relative to the average template. Once this was confirmed, automatic segmentation was carried out to separate the skull from both lower jaws. After that, a suite of image deformation and normalization was used separately for each segmented structure and for all specimens relative to the average atlas. All image transformations generated in the previous step were then applied to propagate the landmark configuration digitized onto the average template onto each segmented structure and for all specimens via function antsApplyTransformsToPoints(). All automatically placed landmarks for each structure were compiled into a single dataset for further geometric morphometric analyses.

### Geometric morphometric analyses

Since the skull has an instance of object symmetry and the lower jaws exhibit matching symmetry, the original skull landmark configuration was duplicated, then the paired landmarks from this copy were swapped (relabelling) while all landmark configurations from the left lower jaw were flipped (by multiplying all x coordinates by-1) to correspond to the landmark configurations digitized on the right lower jaw (Kent and Mardia 2001; Mardia et al. 2000).

A generalized Procrustes analysis (GPA) applied to the full landmark dataset extracted shape by removing extraneous effects of location, orientation, and position (e.g., Dryden and Mardia 1998). After that, the Procrustes coordinates were averaged by individual to extract the component of symmetric shape variation while discarding the asymmetry. A principal component analysis (PCA) was used on the covariance matrix of the Procrustes coordinates to extract the PC scores later used as shape data in subsequent analyses. Centroid size, the most common and explicit measure of size in geometric morphometrics, was computed as the square root of the sum of the squared distances of all landmarks from their centroid (e.g., Slice et al. 1996).

We assessed the effects of age and areal Bone-mineral density (aBMD) since there is a known predisposition of CFW mice toward abnormally high aBMD on shape (Pallares et al. (2015)) by multivariate regressions using 10,000 rounds of permutations separately for the skull and lower jaws as done in the original study. For the skull, there was no association between age and shape (r^2^ = 0.00053, p = 0.75) and a subtle but significant association between aBMD and shape (r^2^ = 0.00561, p = 0.033). For the lower jaws, there were subtle but significant associations between age and shape (r^2^ = 0.00628, p = 0.0285) and between aBMD and shape (r^2^ = 0.00608, p = 0.0283). Given these weak associations, aBMD and age were not used as covariates in the subsequent trait mapping.

All analyses were done with the functions ‘gpagen()’ for the GPA, ‘bilat.symmetry()’ for object symmetry, and ‘procD.lm()’ for the multivariate regressions from the R package geomorph (Adams and Otárola-Castillo 2013).

The average atlas was landmarked twice to test for digitizing error via the traditional two-way mixed model Procrustes ANOVA and ANOVA used in studies of fluctuating asymmetry (FA) (Leamy 1984; Palmer and Strobeck 1986; Savriama et al. 2017; Savriama and Klingenberg 2011). FA refers to small random differences between left and right sides of bilaterally symmetric structures and is, therefore, the smallest level of biological variation to which digitizing error is compared. Analysis was done in MorphoJ (Klingenberg 2011).

### Genome-wide analyses

For QTL mapping, we use the same approach as in the original study. Briefly, PCs summarizing at least 1% of variance were used for a univariate QTL mapping, and centroid size was considered for size QTL mapping. Each PC was analyzed separately.

Genome scans were conducted via the linear mixed model (LMM) approach in GEMMA version 0.98.1 (Zhou and Stephens 2012) while accounting for the relationships among individuals (kinship matrix) and using the ‘leave one chromosome out’ (LOCO) method in which a kinship matrix is calculated using markers from all other chromosomes except the ones that are on the chromosome under consideration (Cheng et al. 2013; Parker et al. 2014). We used the p-value computed from the likelihood ratio test output from GEMMA as the association test statistic. Permutation tests (1000 rounds) were used to define genome-wide significance thresholds separately for each of the phenotypes used in the mapping, as well as an average significance threshold per structure, including only phenotypes for which a significant association was found.

Data from Pallares et al. (2015) were also re-analyzed with the same pipelines for further detailed comparisons between the two landmarking methods. The analysis confirmed the previous findings.

In addition, the function scanoneShape() from the R package shapeQTL (Navarro 2015) was used for multivariate mapping (Maga et al. 2015; Navarro and Maga 2016) using the same PCs included in the univariate mapping to further compare both landmarking methods. Shape is inherently multivariate, hence the use of this approach which simultaneously includes all PCs. Permutation tests (1000 rounds) were used to define single genome-wide significance thresholds separately for each structure.

Here, it was not possible to define QTL regions based on the linkage disequilibrium (LD) pattern around the significant SNPs due to a sparse LD signal for all peak SNPs identified via the automatic landmarking approach (not shown here). Since this sometimes was also the case in the original study, Bayes credible intervals of QTL (Dupuis and Siegmund 1999; Maga et al. 2015; Manichaikul et al. 2006; Navarro and Maga 2016; Sen and Churchill 2001) were instead calculated for the peak SNPs found in both landmarking methods via function bayesint() in the R/qtl package (Broman et al. 2003).

Overlap of relevant QTL regions determined in both landmarking methods was assessed via functions IRanges() and findOverlaps() from the GenomicRanges R package (Lawrence et al. 2013).

The position of peak SNPs identified in the automatic method was visualized via function chromPlot() from the R package of the same name (Oróstica and Verdugo 2016), and overlapping QTL regions were highlighted via the zoom-in method implemented in the function chromoMap() from the R package of the same name (Anand 2019).

## Results

### Digitizing error

The ANOVAs for centroid size and shape for skull and lower jaws both indicate that the ‘Individual-by-Side’ (FA) interaction is highly significant (p < 0.001), which means that the smallest biological variation that can be measured here greatly exceeds the measurement error due to ‘Digitizing’ (Table S2). Note that there is no centroid size asymmetry for the skull since a unique configuration was considered for the whole structure and due to geometric constraints imposed by the GPA.

### QTL mapping and overlap of QTL regions

For the atlas-based method, we kept all PCs summarizing at least 1% of the total variance as in the original study, and this included the first 3 PCs, which together account for more than 99% of the variance of the skull and lower jaws shape. In the original study of Pallares et al. (2015), 22 PCs accounting for 84% of skull shape variation and 21 PCs representing 94% of mandible shape variation were used in the univariate mapping method, and the same PCs were used in the multivariate mapping as well. The average genome-wide significance thresholds used in the univariate mapping was 6 for both structures using the semi-automatic landmark data and 5.78 for the skull, and 6.04 for the lower jaws using the automatic landmark data. This threshold was 5.77 for the skull and 5.80 for the lower jaws using the semi-automatic landmark data and 5.56 for the skull, and 5.50 for the lower jaws using the automatic landmark data in the multivariate mapping.

In the univariate mapping of the skull, none of the significant SNPs of the original study were recovered using the automatic landmarking approach and only one out of the 17 QTLs (Figures 2A, B). The overlap is seen for a QTL around the EGF pathway gene *Gab1* on chromosome 8 for the marker ‘rs36374407′, which has its QTL region overlapping with the one defined for the corresponding marker ‘rs228570244’ found in the original study (Figure 2G). Instead, a new QTL associated with the marker ‘rs231004467’ on chromosome 6 was found with the automatic method (Figures 2A, B).

**Figure 2.**
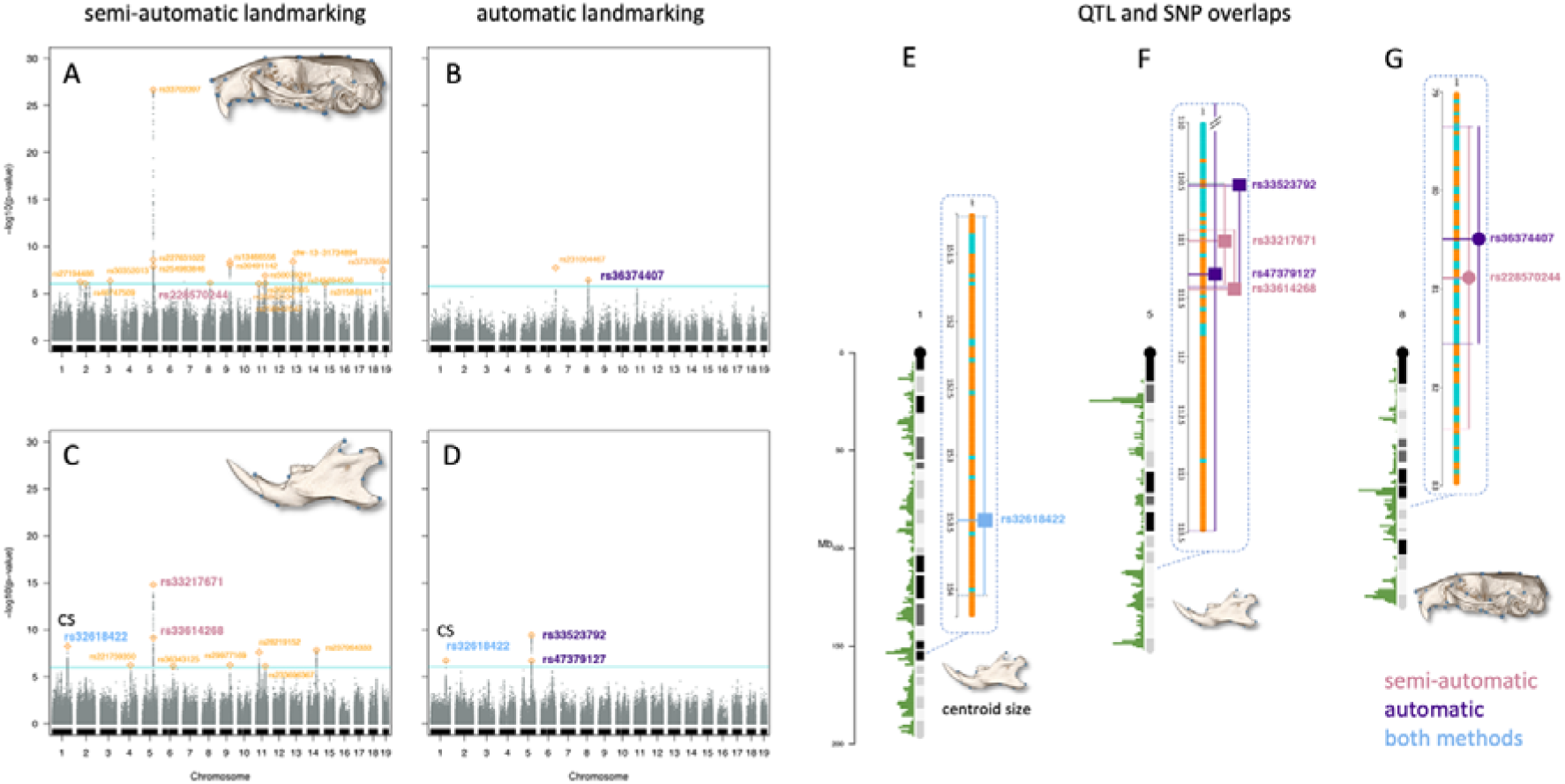
Univariate mapping and locations of SNPs. Genome-wide scans for semi-automatic landmarking with manual adjustment (re-analysis of data from Pallares et al. 2015) and automatic landmarking (this study). (A, B) for the skull and (C, D) for the lower jaws. (E - G) overlapping QTL regions highlighted via zoom-in with 95% Bayesian credible intervals are indicated by lines from either side of each QTL. Orange: non-overlapping QTL regions. Fuchsia and Purple: markers with overlapping QTL regions. Blue: the same marker found in both methods. Square: markers for the lower jaws. Circle: markers for the skull. Marker positions and statistics are provided in Table S3.

For the univariate mapping of the lower jaws, none of the 9 significant SNPs in 8 QTLs of the original study were recovered using the automatic landmarking approach (Figure 2C, D). However, an overlap with two new significant SNPs is seen for a QTL around the transcriptional activator *Mn1* on chromosome 5 for the markers ‘rs33523792’ and ‘rs47379127′, which have their QTL region overlapping within the one defined for the corresponding markers ‘rs33217671’ and ‘rs33614268’ found in the original study (Figure 2F).

For centroid size, no QTL was found for the skull, and one QTL on chromosome 1 was identified for the lower jaws. This is consistent with the results reported in the original study. For the lower jaws, the same peak SNP was recovered by the automatic method (Figure 2E).

In the multivariate mapping, none of the four skull QTLs found with the semi-automatic landmark data were recovered using the automatic landmarking approach, and the only marker ‘cfw−5−46967657’ identified on chromosome 5 does not have its QTL region overlapping with the other marker ‘rs32067860’ defined on the same chromosome with the semi-automatic landmark data (Figures 3A, B). For the lower jaws, two QTLs were discovered using the automatic landmarking approach with the same major SNP ‘rs29385180’ that was also found using the semi-automatic landmark data, which also happened to be the sole QTL found in this case (Figures 3C, D, and E).

**Figure 3.**
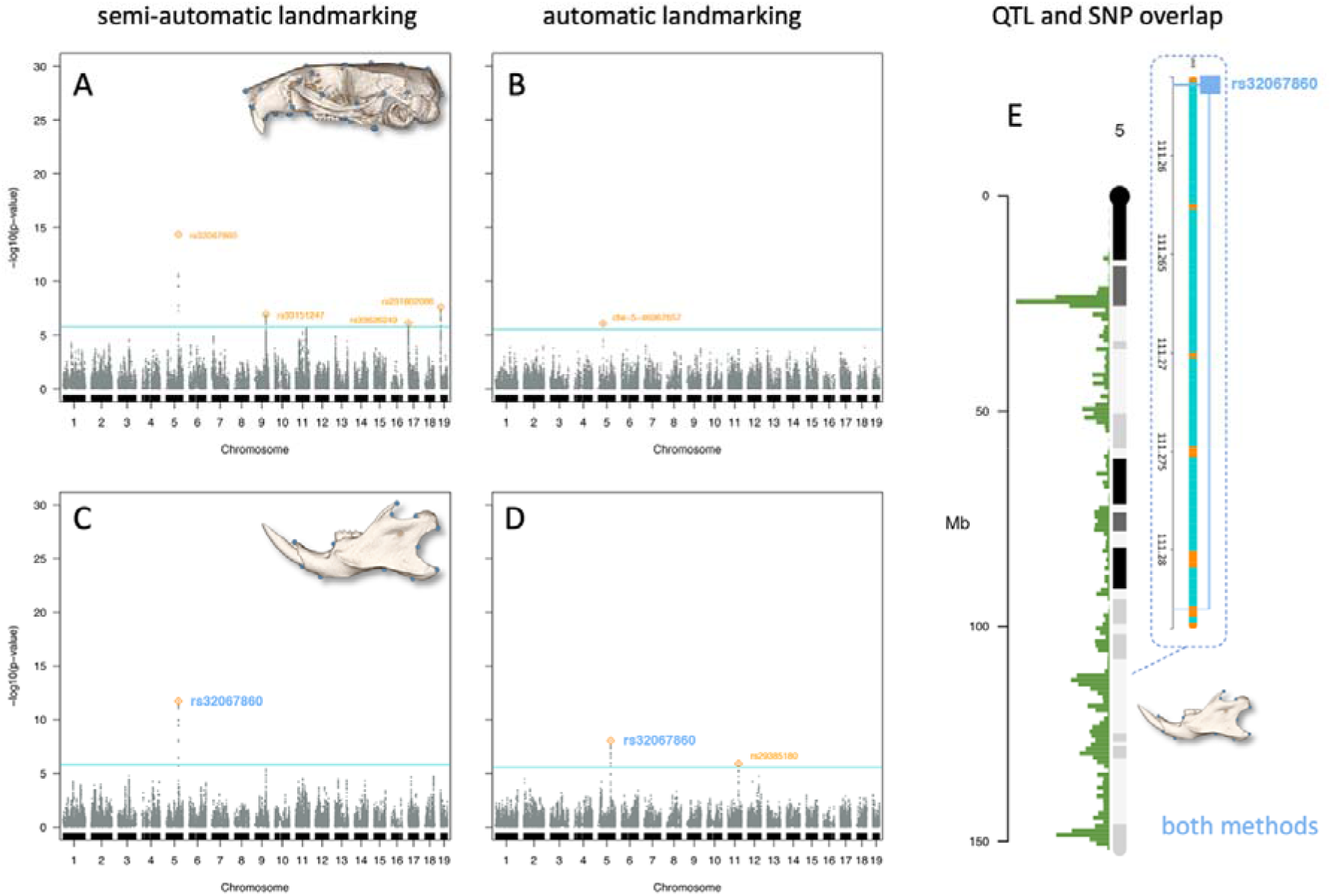
Multivariate mapping and locations of SNPs. Genome-wide scans for semi-automatic landmarking with manual adjustment (re-analysis of data from Pallares et al. 2015) and automatic landmarking (this study). (A, B) for the skull and (C, D) for the lower jaws. (E) overlapping QTL regions highlighted via zoom-in with 95% Bayesian credible intervals are indicated by lines from either side of each QTL. Orange: non-overlapping QTL regions. Blue: the same marker found in both methods. Square: markers for the lower jaws. Circle: markers for the skull. Marker positions and statistics are provided in Table S3.

## Discussion

Our results suggest that the accuracy of the automatic method for detecting variation required for QTL mapping is compromised, regardless of the mapping method used, though somewhat depending on the structure analyzed. For the skull that reflects a structure of a 3D organization, we found that most of the informative variance was lost, given the failure to detect the majority of QTLs from the previous study of Pallares et al. (2015). The informative variance of the lower jaw, which represents an object that is more 2D-like, where its outline can almost fully characterize the shape, was more adequately captured by the automatic method in either mapping technique, but still with less resolution than for the semi-automatic method. For centroid size of the lower jaws, the automatic method recovers the same QTL and the same SNP as the semi-automatic method. These identical results in both methods for this particular trait could be related to the fact that such a univariate measure of size seems easier to capture than a multivariate trait such as shape.

Our results do not appear to concur with the current trend accepting that such automatic method seems to be a reliable and promising tool, despite the known and quite often systematic differences between manual and automatic landmarking often evaluated either by linear distances and analyses of differences between group means and covariance matrices (Maga et al. 2017; Percival et al. 2019; Porto et al. 2020). In these particular studies, despite the known fact that the automatic landmarking procedure reduces the variance given its framework, the biological signal of interest is still captured via this method since shape differences between inbred mouse genotypes are known to be rather large. The atlas-based automatic method that relies on a suite of geometric transformations and image deformations to backpropagate the template’s landmark configuration onto each specimen captures the actual biological signal only to some extent. Given that there is not much difference between mapping techniques, the automatic method causes this departure from the results obtained with the manual approach.

Here, we have illustrated and tested how the known differences between the landmarking approaches affect analyses. Our results suggest strong cautions regarding using such atlas-based automatic landmarking in geometric morphometric analyses, especially in GWAS, where the biological signal of interest is relatively small and cannot be adequately captured by it. Such mapping analyses can, in turn, be considered as an accurate and convenient tool for testing landmarking precision whenever applicable.

## Supporting information

pheno.auto.skull.mand data Table

## Data Availability

The complete code reproducing the analyses in Pallares et al. (2015) as well as in this study and data from Pallares et al. (2015) have been deposited at http://dx.doi.org/10.5061/dryad.k543p. Data generated for this study are available in the supplementary data.

The software used in this study for template building and the R code for automatic segmentation and landmarking are freely available at:

https://github.com/ANTsX/ANTs
https://github.com/ANTsX/ANTsR
https://github.com/muratmaga/mouse_CT_atlas

## Ethics Statement

No new animal experiments have been carried out since we re-used CT-scans that have been already produced in the reference study (Pallares et al. 2015).

## Acknowledgments

We thank Abraham Palmer for allowing the re-analysis of CT scans of mice generated by the MaxPlanck Institute for Evolutionary Biology (Plön, Germany) that were part of a project supported by his National Institute of Health grant (R01GM097737) received from the University of California San Diego. We thank Nicolas Navarro and Murat Maga for their constructive remarks and comments on our work and Sylvain Gerber for stimulating discussions. We also thank Derk Wachsmuth, Kristian Ullrich, and Werner Wegner from our IT services and “Wallace” High Performance Computing Cluster, Carsten Fortmann-Grote from the Scientific Computing Unit at the MPI for Evolutionary Biology, the Max-Planck computing facilities at the Gesellschaft für Wissenschaftliche Datenverarbeitung Göttingen (GWDG), and the Max-Planck Computing and Data Facility (MPCDF).

## Author Contributions

YS and DT conceived the project and designed the analyses. Y.S. processed the CT scans, collected and analyzed the data. YS and DT wrote the manuscript. Both authors read and approved the final manuscript.

## Supplementary material

**Table S1.** Definition of skull and lower jaws landmarks (same as in Pallares et al. 2015)

**Table S2.** Two-way mixed model ANOVA and Procrustes ANOVA for digitizing error.

**Table S3.** Univariate mapping. QTLs identified in this study and overlapping ones found with the original study (re-analysis of data from Pallares et al. 2015). CS: Centroid size, chr: chromosome, pos: position in Mbp, log10p: log10-transformed association P values for each mapping, Lower.CI and Upper.CI: Bayesian estimates of their confidence intervals.

**Supplementary data.** Data generated via the automatic phenotyping method subsequently used in both the univariate and multivariate QTL mapping (file ‘pheno.auto.skull.mand.csv’). This file contains principal component (PC) scores and centroid size (CS) for skull and lower jaws.

**Table S1.**
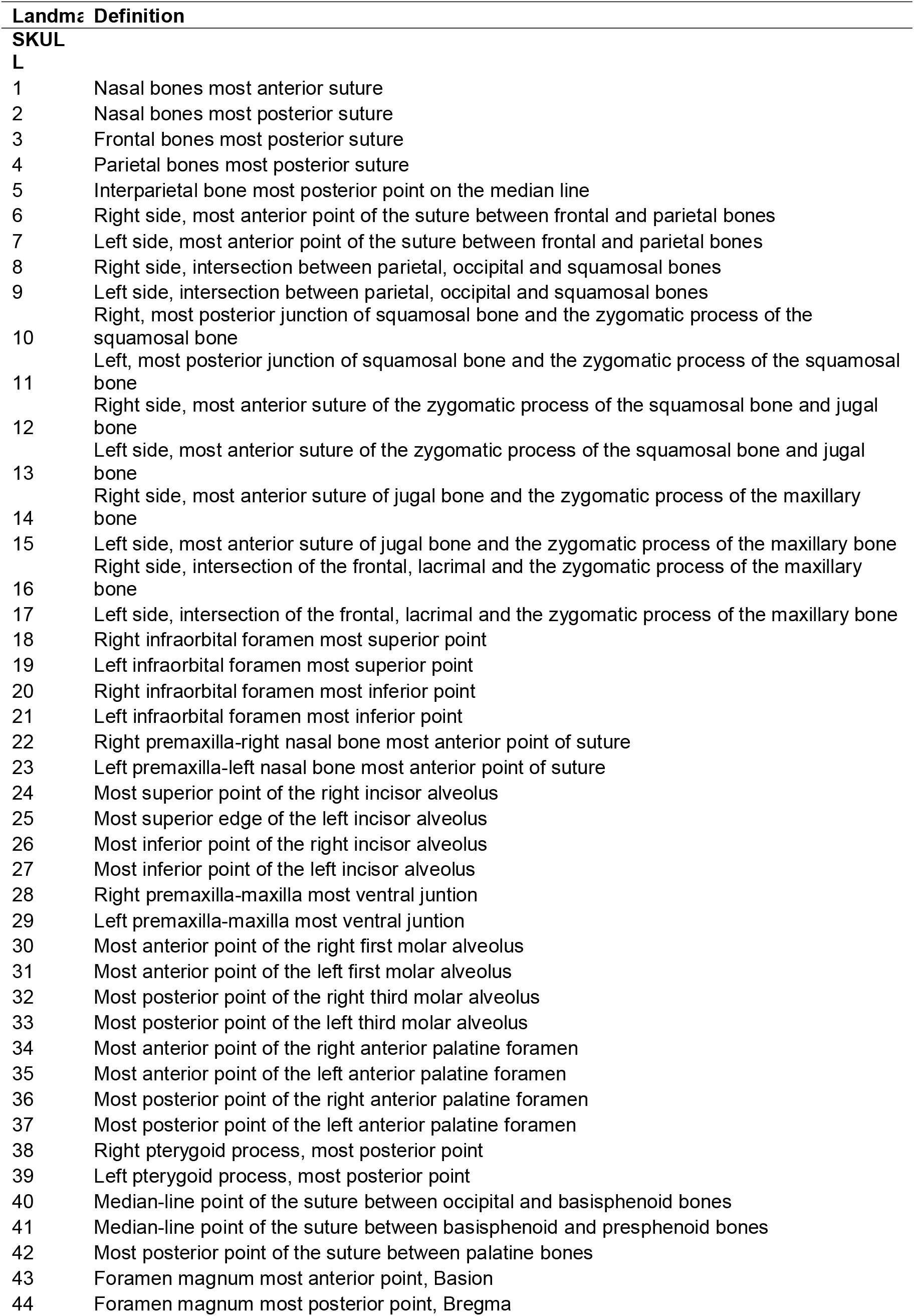
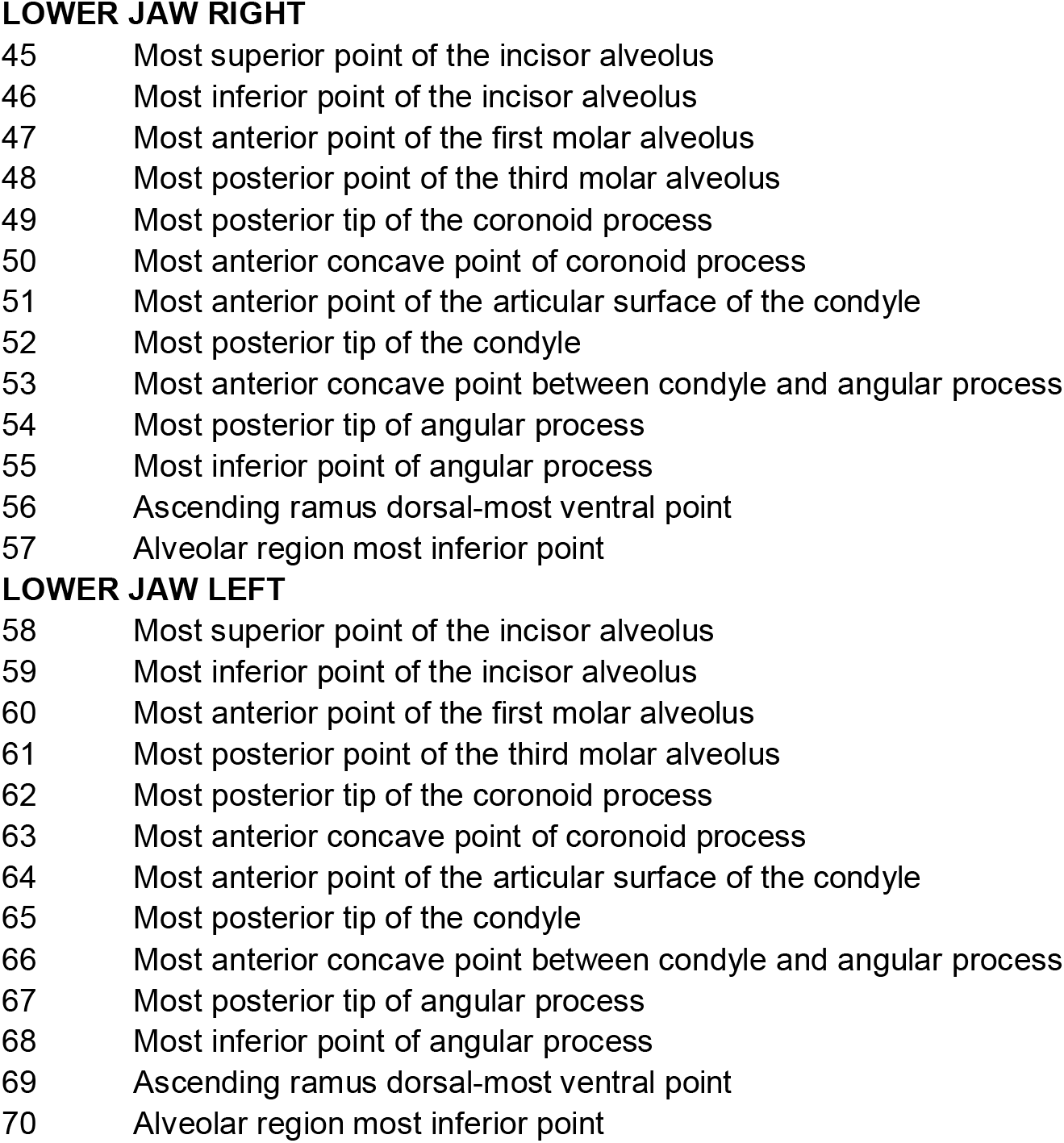
Definition of skull and lower jaws landmarks (same as in Pallares et al. 2015)

**Table S2.** Two-way mixed model ANOVA and Procrustes ANOVA for digitizing error.

Centroid size ANOVA
| **Effect** | **SS** | **MS** | **df** | **F** | **P** |
| --- | --- | --- | --- | --- | --- |
| Individual | 789.415447 | 1.145741 | 689 | 1805.24 | <.0001 |
| DigitizingError | 0.437926 | 0.000635 | 690 |  |  |

Procrustes ANOVA
| **Effect** | **SS** | **MS** | **df** | **F** | **P** |
| --- | --- | --- | --- | --- | --- |
| Individual | 0.12413394 | 2.689E-06 | 46163 | 9.06 | <.0001 |
| Side | 0.18621415 | 0.00321059 | 58 | 10814.33 | <.0001 |
| Ind*Side | 0.01186403 | 2.969E-07 | 39962 | 1.42 | <.0001 |
| DigitizingError | 0.01805619 | 2.093E-07 | 86250 |  |  |

**Lower Jaws** Centroid size ANOVA
| **Effect** | **SS** | **MS** | **df** | **F** | **P** |
| --- | --- | --- | --- | --- | --- |
| Individual | 1393.69134 | 2.022774 | 689 | 1.31 | 0.0002 |
| Side | 1.639829 | 1.639829 | 1 | 1.07 | 0.3024 |
| Ind*Side | 1060.8179 | 1.539649 | 689 | 15474.81 | <.0001 |
| DigitizingError | 0.137302 | 0.000099 | 1380 |  |  |

Procrustes ANOVA
| **Effect** | **SS** | **MS** | **df** | **F** | **P** |
| --- | --- | --- | --- | --- | --- |
| Individual | 1.19876706 | 5.4371E-05 | 22048 | 1.18 | <.0001 |
| Side | 1.28276514 | 0.04008641 | 32 | 866.63 | <.0001 |
| Ind*Side | 1.0198361 | 4.6255E-05 | 22048 | 23.27 | <.0001 |
| DigitizingError | 0.08778533 | 1.9879E-06 | 44160 |  |  |

**Table S3.** Univariate mapping. QTLs identified in this study and overlapping ones found with the original study (re-analysis of data from Pallares et al. 2015). CS: Centroid size, chr: chromosome, pos: position in Mbp, log10p: log10-transformed association P values for each mapping, Lower.CI and Upper.CI: Bayesian estimates of their confidence intervals.

| landmarking method | structure | trait | snp | chr | pos | log10p | Lower.CI | Upper.CI | overlap |
| --- | --- | --- | --- | --- | --- | --- | --- | --- | --- |
| Automatic (this study) | lower jaws | centroid size | rs32618422* | 1 | 153481175 | 6.6928027225081<br>8 | 151223012 | 154041054 | TRUE |
| Automatic (this study) | lower jaws | PC2 | rs33523792 | 5 | 110540748 | 9.3950246254860<br>3 | 110540748 | 111394692 | TRUE |
| Automatic (this study) | lower jaws | PC3 | rs47379127 | 5 | 111298758 | 6.6792469568504<br>3 | 30919182 | 113493026 | TRUE |
| Automatic (this study) | skull | PC1 | rs231004467 | 6 | 147957851 | 7.7170357063819<br>5 | 147957851 | 148393505 | FALSE |
| Automatic (this study) | skull | PC1 | rs36374407 | 8 | 80493535 | 6.4056361825254<br>9 | 79347918 | 81565134 | TRUE |
| Semi-Automatic with manual adjustement (Pallares et al. 2015) | lower jaws | centroid size | rs32618422* | 1 | 153481175 | 8.2247792448792<br>3 | 151223012 | 154041054 | TRUE |
| Semi-Automatic with manual adjustement (Pallares et al. 2015) | lower jaws | PC4 | rs33217671 | 5 | 111018365 | 14.780732797537<br>6 | 110542538 | 111394692 | TRUE |
| Semi-Automatic with manual adjustement (Pallares et al. 2015) | lower jaws | PC7 | rs33614268 | 5 | 111426493 | 9.1176764337703<br>6 | 110918274 | 111426493 | TRUE |
| Semi-Automatic with manual adjustement (Pallares et al. 2015) | skull | PC2 | rs228570244 | 8 | 80889309 | 6.1228656800062<br>4 | 79348411 | 82427354 | TRUE |

| landmarking method | structure | trait | snp | chr | pos | log10p | Lower.CI | Upper.CI | overlap |
| --- | --- | --- | --- | --- | --- | --- | --- | --- | --- |
| Automatic (this study) | lower jaws | shape | rs29385180 | 11 | 94989096 | 5.9056805607284<br>8 | 94932372 | 94989115 | FALSE |
| Automatic (this study) | lower jaws | shape | rs32067860* | 5 | 111256383 | 8.0349453008641<br>6 | 111256383 | 111283008 | TRUE |
| Automatic (this study) | skull | shape | cfw-5-46967657 | 5 | 46967657 | 6.1078942873562<br>3 | 46930846 | 46974028 | FALSE |
| Semi-Automatic with manual adjustement (Pallares et al. 2015) | skull | shape | rs32067860 | 5 | 111256383 | 14.373183118806<br>3 | 111256383 | 111256383 | FALSE |
| Semi-Automatic with manual adjustement (Pallares et al. 2015) | skull | shape | rs33151247 | 9 | 97861578 | 6.9295566980095<br>6 | 97643517 | 99668200 | FALSE |
| Semi-Automatic with manual adjustement (Pallares et al. 2015) | skull | shape | rs33626249 | 17 | 4523620 | 6.1419473722968<br>5 | 3354573 | 4605201 | FALSE |
| Semi-Automatic with manual adjustement (Pallares et al. 2015) | skull | shape | rs231802086 | 19 | 4454653 | 7.6395238452858<br>5 | 4182181 | 4855351 | FALSE |
| Semi-Automatic with manual adjustement (Pallares et al. 2015) | lower jaws | shape | rs32067860* | 5 | 111256383 | 11.711697113912<br>2 | 111256383 | 111283008 | TRUE |

